# Age Dependent Immunopathology Drives Pneumonia-Associated Cardiac Dysfunction During *Streptococcus pneumoniae* Infection

**DOI:** 10.64898/2026.05.19.726323

**Authors:** Sameer Salam Matoo, Naresh Kumar, Mariam A. Salem, Alison T. Thomas, Aakash Shah, Pankaj Kumar, Miqdad O. Dhariwala, Liao Zhiwei, Haitao Wen, Stephanie Sevaue, Latha P. Ganesan, William P. Lafuse, Rachael Philips, Patrick L. Collins, Murugesan V. S. Rajaram

**Affiliations:** Department of Microbial Infection and Immunity, The Ohio State University, Columbus, OH, 43210; Department of Internal Medicine, The Ohio State University, Columbus, OH, 43210; Department of Dermatology, College of Medicine, The Ohio State University, Columbus, OH, 43210

## Abstract

Aging is a major risk factor for severe *Streptococcus pneumoniae* (*Spn*) infection and pneumonia-associated – major adverse cardiac events (PA-MACE), yet underlying mechanisms remain unclear. Using young and aged murine models, we show that aging exacerbates bacterial burden, mortality, and cardiac dysfunction following *Spn* infection, associated with impaired macrophage bacterial killing. Single-cell RNA sequencing infected hearts revealed extensive age-dependent remodeling across immune and stromal compartments. Aged mice exhibited heightened pro-inflammatory myeloid responses, with increased neutrophil infiltration characterized by elevated S100A8/9 and LCN2 and reduced antimicrobial programs. Macrophages displayed defective efferocytosis, including disruption of the GAS6–AXL axis. Aging also drove expansion of an infection-responsive fibroblast population with inflammatory signatures and reduced extracellular matrix gene expression. These changes were linked to oxidative stress and impaired glucose oxidation. Notably, anti-inflammatory treatment rescued cardiac dysfunction, implicating excessive inflammation as a central driver of PA-MACE. Together, these findings define mechanisms linking aging to pneumococcal cardiac complications and identify potential therapeutic targets.

## Introduction

*Streptococcus pneumoniae* (pneumococcus) remains a leading cause of community-acquired pneumonia (CAP) and invasive pneumococcal disease (IPD), contributing substantially to global morbidity and mortality^1–3^. Older adults bear a disproportionate burden of severe disease, hospitalization, and mortality. Despite the availability of vaccines and antibiotic therapies, clinical outcomes in the elderly remain poor, highlighting persistent age-associated defects in host immune defense^2,4^.

Beyond the lung, *S. pneumoniae* (*Spn*) can disseminate systemically and invade the heart, where it induces myocardial injury, arrhythmias, and long-term cardiac dysfunction^5,6^, collectively termed pneumonia-associated major adverse cardiac events (PA-MACE)^7^. These complications markedly increase both short-term mortality and long-term cardiovascular risk^8^. Notably, the risk of heart failure persists for years following infection, and experimental models recapitulate sustained cardiac dysfunction even after antibiotic-mediated bacterial clearance^9–11910,11^. Approximately one in five individuals hospitalized for pneumococcal pneumonia experience PA-MACE, with a fourfold increase in mortality^3^. This burden is particularly pronounced in older adults, yet the mechanisms linking aging to pneumococcus-induced cardiac dysfunction remain poorly defined. Innate immune cells, particularly macrophages and neutrophils, are central to pneumococcal clearance^12^. However, aging is associated with profound immune remodeling, including impaired phagocytosis, reduced microbial killing, dysregulated cytokine production, and chronic low-grade inflammation (“inflammaging”)^13–16^. These changes compromise pathogen control while amplifying inflammatory tissue damage. In parallel, cardiac aging is characterized by fibrosis, extracellular matrix remodeling, endothelial dysfunction, and dysregulated stromal cell activation^17–19^. How these immune and stromal alterations intersect during pneumococcal infection to drive cardiac injury remains unclear. Given the rapid expansion of the global aging population, the burden of pneumococcal pneumonia and its complications is expected to rise substantially. According to projections by the World Health Organization (WHO), by 2030, one in six people worldwide will be aged ≥ 65 years^20^, and in the United States, older adults, particularly those with comorbidities, experience disproportionately high rates of pneumococcal disease and hospitalization ^21,22^. Defining how aging reshapes host responses to infection is therefore critical for developing targeted strategies to mitigate cardiac complications.

In the present study, we investigated how aging alters cardiac immune and stromal responses during pneumococcal infection. Using young and aged murine models, we demonstrate that aging exacerbates cardiac dysfunction and inflammation following *Spn* infection. Through single-cell RNA sequencing and complementary validation, we identify age-dependent reprogramming of myeloid and fibroblast populations, characterized by heightened inflammatory signaling and impaired effector functions, including defective degranulation and efferocytosis. Notably, neutrophils persist in the aged heart during convalescence, and a distinct fibroblast subset emerges selectively in aged, infected tissue. Together, these findings reveal coordinated immune–stromal dysregulation as a key driver of cardiac pathology in aging and provide mechanistic insight into the heightened vulnerability of older adults to pneumococcal disease.

## Results

### Aged mice are highly susceptible to *Spn* infection

Pneumonia mediated invasive pneumococcal disease (IPD) is a major driver of multi-organ dysfunction syndrome, heart failure and increased mortality^36,41,42^ .To isolate pulmonary immune responses, we used an intraperitoneal (IP) infection model of invasive pneumococcal disease (IPD). To evaluate age-associated differences in host susceptibility, young (10-12-week-old) and aged (18–20-month-old) C57BL/6 mice were infected intraperitoneally with 0.5×10³ CFU *Spn*, strain TIGR4 to model IPD. Both groups received ampicillin (100 mg/kg) beginning at 28-hours post-infection (hpi) and every 10 hours thereafter for a total of four doses (**Fig.1A**). Aged mice exhibited markedly reduced survival, with 80% mortality rate compared to 20% mortality in young mice at 96-hpi (**Fig.1B**), indicating pronounced age-related vulnerability. Consistently, aged mice developed significantly higher circulating bacterial burdens (**Fig.1C**). We observed similar findings following intranasal infection, with aged mice exhibiting increased susceptibility and pronounced cardiac dysfunction during *Spn* infection **(Supplementary Fig. 1A–D)**. It has been demonstrated that the release of pneumococcal toxin pneumolysin during bacterial lysis has been implicated in host tissue injury.^23^ To determine whether antibiotic-mediated bacterial lysis and pneumolysin release contributed to mortality, aged mice were treated with either bactericidal ampicillin or the bacteriostatic agent azithromycin (100 mg /kg) beginning at 28 hpi. Bacteriostatic treatment did not improve survival (**Supplementary Fig.2**), suggesting that increased bacterial lysis is not the primary driver of disease severity in aged mice.

**Fig. 1.**
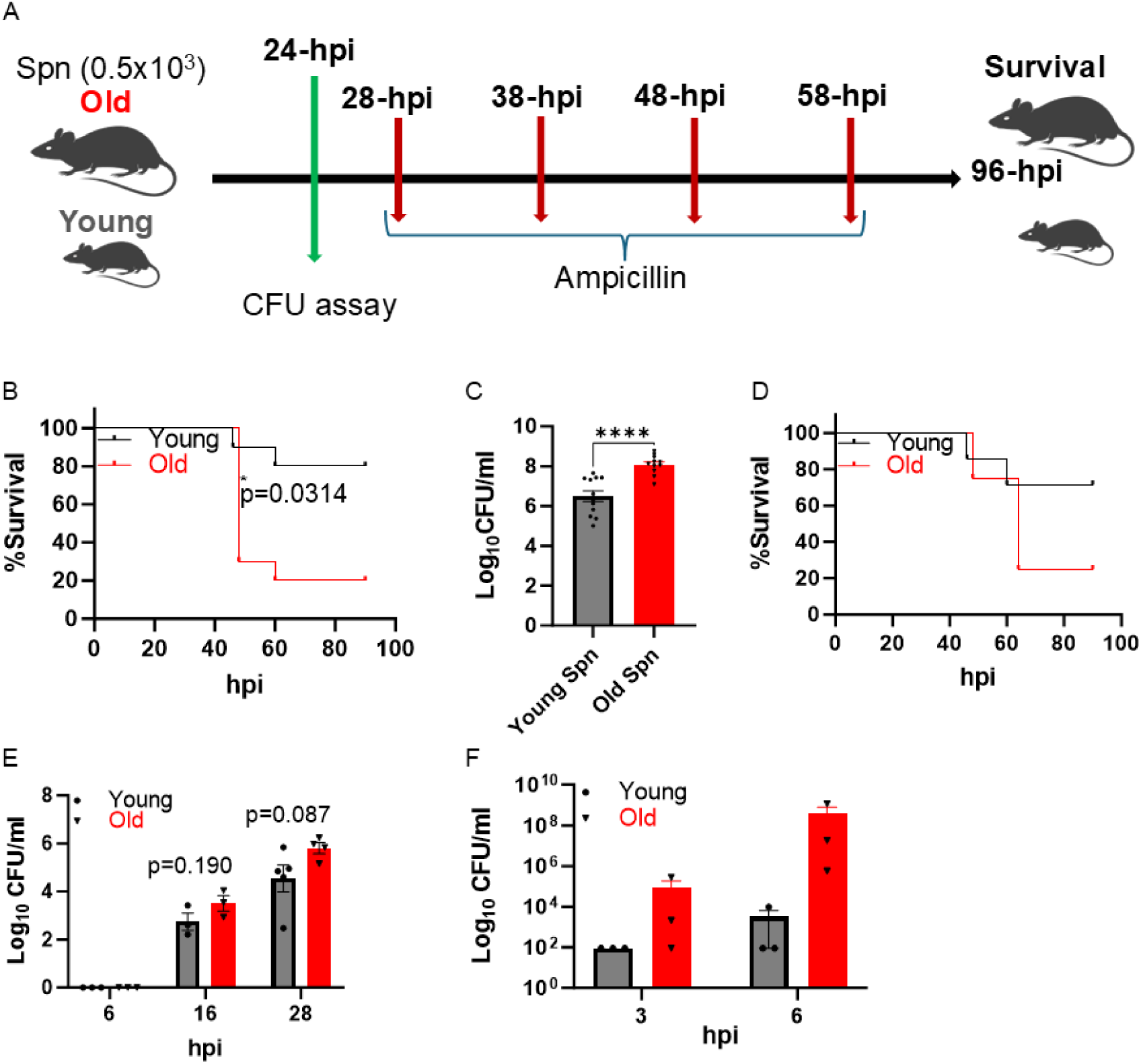
Survival and bacteremia in young and old mice following S. pneumoniae infection. **A)** Schematic of the mouse infection protocol and timeline for data acquisition. **B**) Young and aged C57BL/6 mice were infected intraperitoneally with *Spn*, (0.5×10^3^ CFU); antibiotic treatment was initiated at 28 h post-infection (hpi) and continued every 10 h for 3 days. Survival of young and aged mice is shown (n = 10 per group; experiment performed twice, representative data shown). **C**) Bacterial burden in blood at 24-hpi in young (n=12) and aged (n=11) mice, data pooled from two independent experiments. Young and aged mice were infected with *S. pneumoniae* at age-adjusted doses (young: 0.5 × 10³ CFU; aged: 0.025 × 10³ CFU) and treated with antibiotics as in B. **D**) Survival of young (n = 7) and aged (n = 4) mice (single experiment). **E**) Bacterial burden in blood at 6 and 28 hpi in young and aged mice. **F**) Intracellular bacterial killing by young and aged alveolar macrophages at 3-and 6-hpi. Alveolar macrophages from 3-4 mice were pooled together. L.O.D, limit of detection. Each point represents an individual mouse; bars indicate mean ± SEM. Survival curves were analyzed by log-rank test, and bacterial burden by Student’s t-test. Exact P values are indicated in the panels.

We next evaluated susceptibility at a reduced infectious dose. Aged mice were infected with a 20-fold lower inoculum (0.025×10³ CFU) than that administered to young mice (0.5×10³ CFUs). Remarkably, survival rate of aged, infected mice remained poor, with only 20% surviving at 96-hpi (**Fig.1D**), and displayed bacteremia at 16-and 28-hpi comparable to young mice infected with higher dose of 0.5×10³ CFU (**Fig.1E**). These findings indicate an intrinsic defect in bacterial control with aging. Given the central role of macrophages in early pneumococcal clearance, we assessed macrophage function ex vivo. Alveolar macrophages from aged mice showed comparable bacterial uptake but significantly impaired intracellular killing following infection (MOI = 5 (**Fig.1F**), consistent with prior report^24^. Together, these data identify a macrophage-intrinsic defect in antimicrobial activity that likely contributes to increased bacterial burden and mortality in aged hosts.

### Aged mice exhibit exacerbated cardiac dysfunction during acute infection

To determine whether aging influences cardiac function during acute *Spn* infection, young and aged mice were infected with *Spn* (0.5×10³ CFU) and assessed echocardiography at 24-hpi. The representative cardiac patch shown in Fig. 2A summarizes findings from two independent experiments. Baseline cardiac function was comparable between groups. However, during the acute phase, aged mice developed significantly greater cardiac dysfunction than young, infected mice. Specifically, aged-infected mice exhibited reduced heart rate, stroke volume, ejection fraction and fractional shortening, accompanied by impaired cardiac output (**Fig.2B-F**). These functional deficits coincided with the elevated bacterial burden observed in aged mice (**Fig.1C**). Together, these findings demonstrate that aging exacerbates cardiac dysfunction during the acute phase of pneumococcal infection, indicating increased cardiovascular vulnerability in the aged host under infectious stress.

**Fig. 2.**
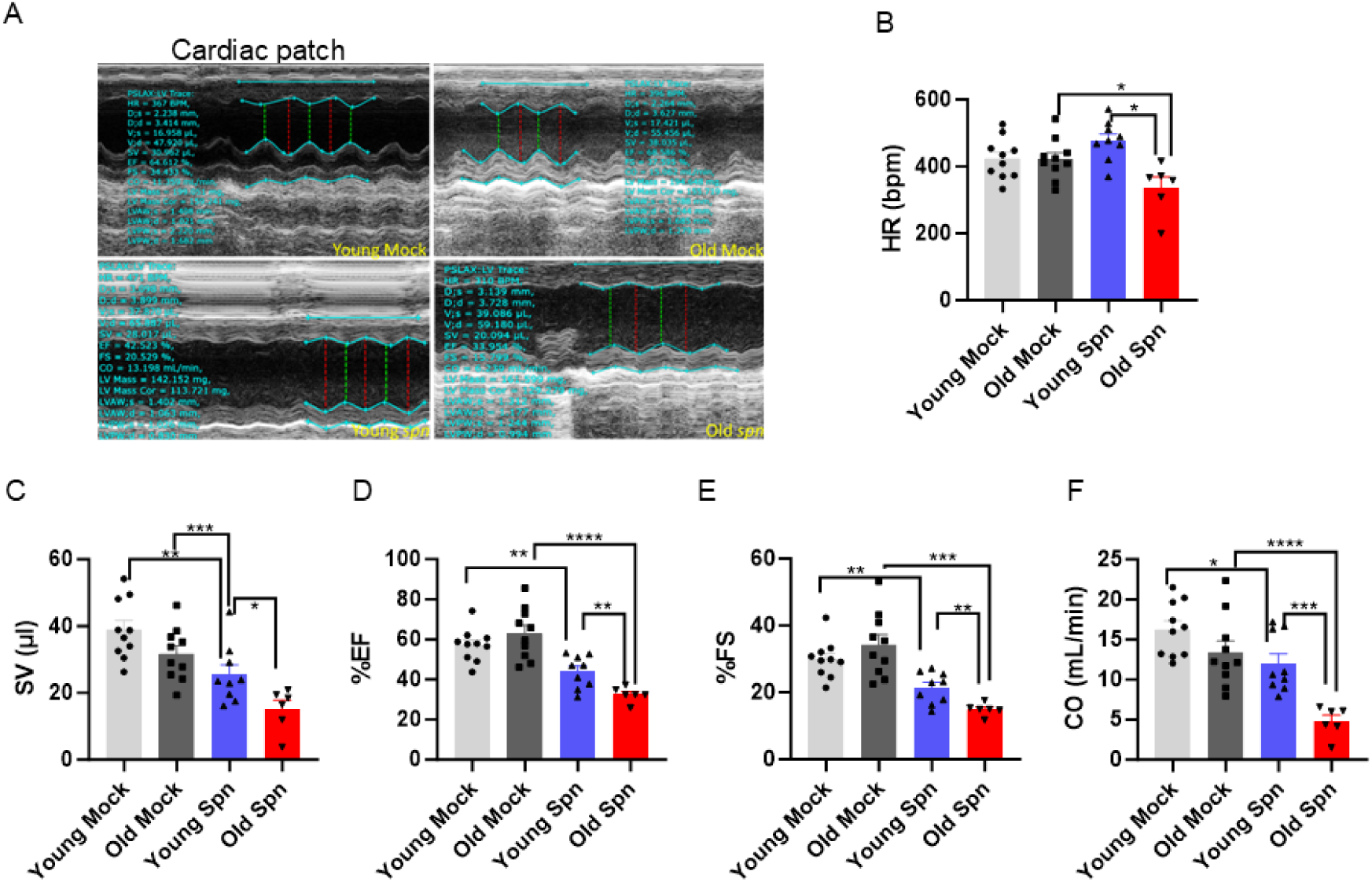
Aged mice exhibit greater cardiac dysfunction during the acute phase of *Spn* infection. Young and aged mice were infected intraperitoneally with *S. pneumoniae* (0.5 × 10³ CFU), and cardiac function was assessed by echocardiography at 24 h post-infection (young mock, n = 10; aged mock, n = 10; young *Spn*, n = 9; aged *Spn*, n = 6). **A**) Representative mode echocardiography images (“cardiac patch”) from young mock, aged mock, young infected, and aged infected mice. **B–F**) Quantification of cardiac parameters, including heart rate (HR; **B**), stroke volume (SV; **C**), ejection fraction (%EF; **D**), fractional shortening (%FS; **E**), and cardiac output (CO; **F**). Each point represents an individual mouse; bars indicate mean ± SEM. Data are pooled from two independent experiments. Statistical comparisons were performed using Student’s test (*P < 0.05, **P < 0.01, *** P < 0.001, ****P < 0.0001).

### Pneumolysin is a key determinant of mortality in aged mice

*Spn* produces multiple virulent factors that contribute to host tissue injury and mortality. Among these, pneumolysin (Ply), a cholesterol-dependent pore-forming toxin, induces direct cardiomyocyte injury, whereas pyruvate oxidase (SpxB) generates hydrogen peroxide (H₂O₂) and promotes oxidative stress^25^. Although both factors have been implicated in pneumococcal pathogenesis, their relative contributions to mortality in aging remain unclear. To address this, aged mice were infected with wild-type (*Spn*), a Ply-deficient strain (Δply), or a strain lacking SpxB (ΔspxB). Infection with wild-type bacteria resulted in rapid mortality, with most aged mice succumbing within the first 24–48 hpi (**Supplementary Fig.3A**). In contrast, Δply-infection significantly improved survival, whereas deletion of SpxB had no protective effect. Notably circulating bacterial burdens at 24 hpi were comparable across groups (**Supplementary Fig.3B**), indicating that Ply drives mortality independently of systemic bacterial load.

Given the well-established association between severe pneumonia and cardiac complications by inducing cardiomyocyte necrosis^23^, we next tested whether *Spn* directly impairs cardiomyocyte contractile function. We used human induced pluripotent stem cell-derived cardiomyocytes (hiPSC-CMs) and multi-electrode array (MEA) system to monitor cardiomyocyte electrical activity. Briefly, we infected hiPSC-CMs were exposed to *Spn* (MOI = 1), and the conduction function was recorded real time for extended period. Infection caused progressive reductions in beating rate, field potential duration, and spike amplitude beginning at 12–16 hpi, whereas mock-treated cells remained stable (**Supplementary Fig.4A–D**). Electrophysiological recordings revealed increasing conduction instability over time, consistent with infection-induced dysfunction (**Supplementary Fig.4E–F**).

Together, these findings identify pneumolysin as a critical mediator of mortality in aged hosts and demonstrate that toxin-driven cardiomyocyte dysfunction, rather than bacterial burden alone, underlies severe disease in aging.

### Single cell-RNA sequencing (scRNA-seq) of cardiac cells revealed cellular heterogeneity during infection

Aging is associated with chronic low-grade inflammation, called inflammaging,^26^ which can fundamentally alter tissue immune tone and stress responses. To investigate how *Spn* infection reshapes cardiac immune and non-immune compartments during the resolution phase, we performed single-cell RNA sequencing (scRNA-seq) on hearts from young and aged mock-infected) and from *Spn*-infected young and aged mice (n=4 per group), dissociated enzymatically, and processed for droplet-based scRNA-seq (**Fig.3A**). This yielded 93,962 high quality cells, with an average of ∼23,000 cells per condition. Unsupervised clustering and annotation based on canonical marker genes revealed robust cellular heterogeneity, including B cells (CD19), endothelial cells (PECAM1), fibroblasts (PDGFRA, COL1A1), lymphatic endothelial cells (PECAM1, PROX1, FLT4), mast cells (KIT, FCER1A), mesothelial cells (MSLN), myeloid cells (CD14, MAFB), neuronal cells (NCAM1), NK/T cells (PTPRC,CD3E), pericytes (PDGFRB), and smooth muscle cells (CNN1, ACTA2) (**Fig.3B,C**). Pearson correlation analysis grouped transcriptionally similar clusters into major immune lineages, further supporting the accuracy of cell-type assignments (**Fig.3D**). The overall composition and marker profiles were consistent with previously reported murine cardiac scRNA-seq atlases, validating dataset quality and providing a detailed framework for dissecting age-and infection-associated remodeling of the cardiac cellular landscape^27^.

**Fig. 3.**
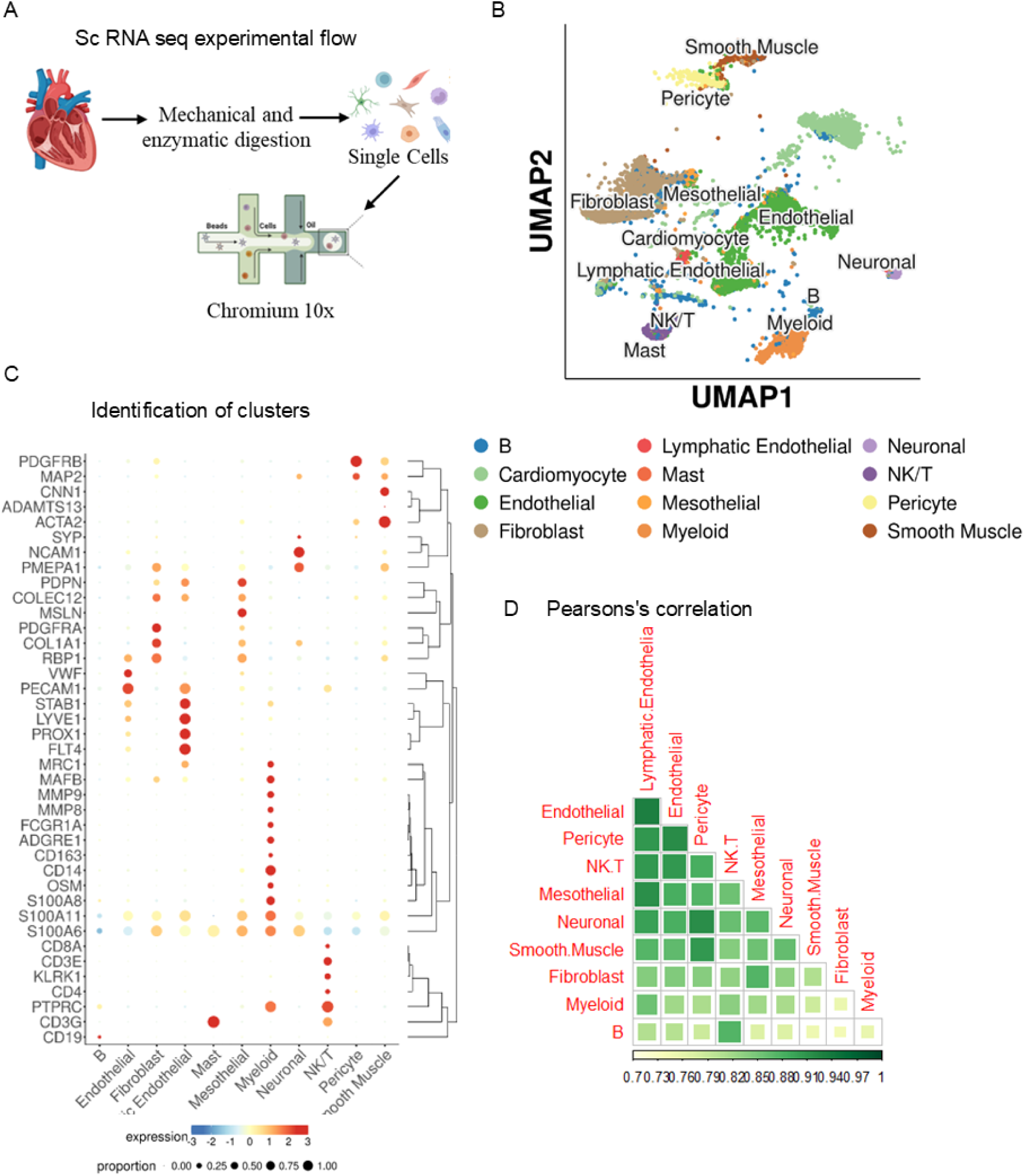
Single-cell RNA sequencing of cardiac tissue reveals distinct cellular clusters following infection. Young and aged mice were infected intraperitoneally with *S. pneumoniae* (0.5 × 10³ CFU), treated with antibiotics starting at 28 h post-infection (hpi) every 10 h for 3 days, and hearts were collected at 96 hpi for single-cell RNA sequencing (young mock, n = 8; aged mock, n = 8; young *Spn*, n = 8; aged *Spn*, n = 2; samples pooled within groups). **A**) Schematic overview of the single-cell RNA-seq workflow used to profile cardiac cells from young and aged mice under mock or *Spn*-infected conditions. **B**) UMAP embedding and clustering of cardiac cells showing transcriptionally distinct populations identified using Seurat-based annotation; some populations, notably cardiomyocytes, did not meet transcriptome quality thresholds for downstream analysis. **C**) Dot plot showing expression of selected marker genes used to define major cardiac cell types. **D**) Pearson correlation heat map illustrating transcriptomic similarity among clusters, supporting cell-type identity and biological relatedness across samples.

### Single-cell profiling reveals age-associated myeloid dysfunction in infected hearts

Unsupervised dimensionality reduction (UMAP) of cardiac myeloid cells identified five clusters, corresponding to neutrophils, macrophage-tissue resident (macrophage-TR), macrophage cycling, monocyte/macrophage, and a distinct myeloid population (**Fig.4A**). All clusters were present across young mock, old mock, young *Spn*, and old *Spn* groups, with macrophage-TR and neutrophils representing the dominant myeloid populations in mock and infected hearts, respectively (**Fig.4B, C**). Each cluster exhibited a distinct transcriptional program, and canonical marker expression confirmed high-resolution delineation of myeloid subtypes (**Supplementary Fig.5A, B**).

**Fig. 4.**
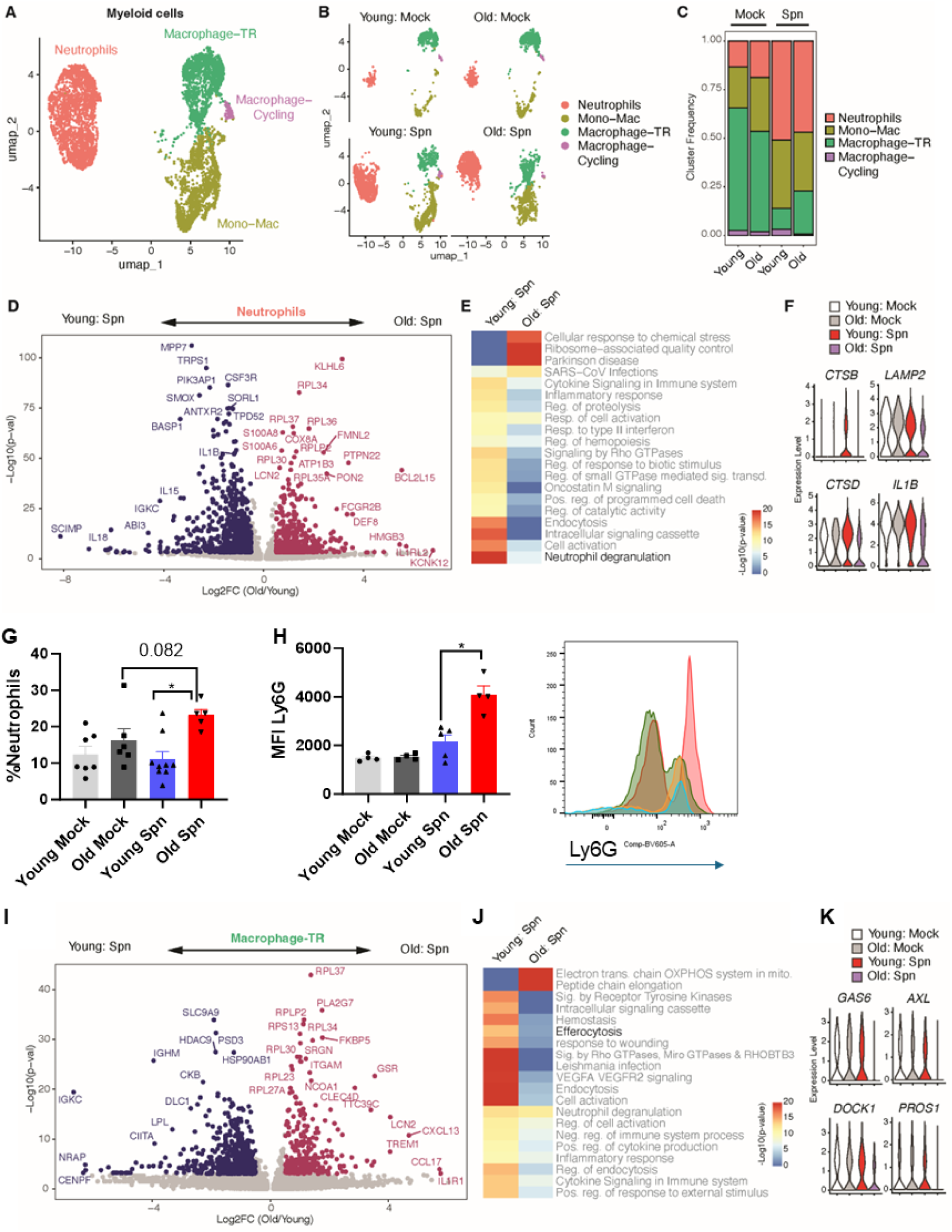
Single-cell profiling reveals age-associated neutrophil and macrophage dysfunction in infected hearts. **A, B**) UMAP visualization and cluster annotation of cardiac myeloid cell populations identified by scRNA-seq at 96-hpi. **C**) Frequencies of myeloid cell clusters across young mock, aged mock, young *Spn*, and aged *Spn* groups. **D**) Differential gene expression analysis of neutrophils from aged versus young *Spn*-infected mice, shown as a volcano plot. **E**) Pathway enrichment analysis in neutrophils from young and aged *Spn*-infected mice. **F**) Violin plots showing expression of neutrophil degranulation- and lysosome-associated markers in neutrophils from young and aged mock- and *Spn*-infected mice. **G**) Quantification of cardiac neutrophil abundance at 96-hpi. **H**) Geometric mean fluorescence intensity (gMFI) of Ly6G on circulating blood neutrophils at 24 hpi. **I**) Differential gene expression analysis of tissue-resident macrophages (Macrophage-TR) from aged versus young *Spn*-infected mice shown as by volcano plot. **J**) Pathway enrichment analysis of tissue-resident macrophages from young and aged *Spn*-infected mice. **K**) Violin plots showing expression of efferocytosis-associated markers in macrophages from young and aged mock- and *Spn*-infected mice. P-values compare young and old mice. ***p < 0.001 by Wilcoxon rank-sum test.

To define age-associated changes in cardiac neutrophils, we compared neutrophils from young and old *Spn*-infected hearts. Differential expression analysis revealed a robust age-dependent transcription (**Fig.4D**). Neutrophils from old, infected hearts upregulated of inflammatory and activation-associated genes, including S100A8, LCN2, FCGR2B, and DEF8, as well as genes linked to metabolic and cytoskeletal remodeling (COX8A, ATP1B3, and FMNL2). In contrast, genes associated with homeostatic and receptor signaling pathways (MPP7, CSF3R, PIK3AP1, and IL15) were downregulated, suggesting impaired regulatory signaling. Pathway enrichment analysis indicated activation of cellular stress, inflammatory, and cytokine-mediated signaling pathways, whereas key neutrophil functions, including endocytosis and degranulation, were attenuated (**Fig.4E**). Consistent with these findings, lysosomal genes such as CTSB, LAMP2, and CTSD were reduced in neutrophils from old, infected hearts (**Fig.4F**), indicating defective degranulation and proteolytic capacity. Flow cytometry confirmed increased neutrophil accumulation in hearts of aged *Spn* -infected mice compared with young counterparts (**Fig.4G**), and aged mice also exhibited elevated. Ly6G geometrics means fluorescence intensity (gMFI)in circulating neutrophils at 24 hpi (**Fig.4H**), consistent with heightened activation and trafficking^28^.

Cardiac resident macrophages play critical roles in inflammation, tissue repair, and resolution^23,45,46^, and we found a reduction in cardiac macrophage frequencies in infected hearts (**Fig.4C**), we next examined age-associated transcriptional changes in macrophages-TR. Macrophages from old *Spn*-infected hearts showed profound reprogramming, with upregulation of genes involved in inflammatory signaling and cellular stress, including TREM1, CCL17, CXCL13, IL1R1, GSR, and ITGAM (**Fig.4I**). Pathway enrichment analysis revealed dysregulation of oxidative phosphorylation, intracellular signaling, VEGF/VEGFR signaling, efferocytosis, and cytokine signaling in aged cardiac macrophages, indicative of metabolic stress and impaired resolution programs (**Fig.4J**). Notably, expression of efferocytosis- and repair-associated genes, including GAS6, AXL, DOCK1, and PROS1, was reduced in aged hearts (**Fig.4K**), suggesting compromised clearance of apoptotic cells and defective termination of inflammation.

### Aging skews cardiac fibroblasts toward inflammatory and maladaptive remodeling phenotypes

Single-cell transcriptomic profiling revealed extensive remodeling of cardiac fibroblast states following *Spn* infection, with aging markedly altering both the magnitude and complexity of this response (**Fig.5**). Unsupervised clustering identified five transcriptionally distinct fibroblast populations: homeostatic-ECM, infection-responsive fibroblasts (InRFs), myofibroblasts, inflammatory fibroblasts, and cycling cells representing discrete functional states within a highly heterogeneous fibroblast compartment (**Fig.5A-C** and **Supplementary Fig.6**).

**Fig. 5.**
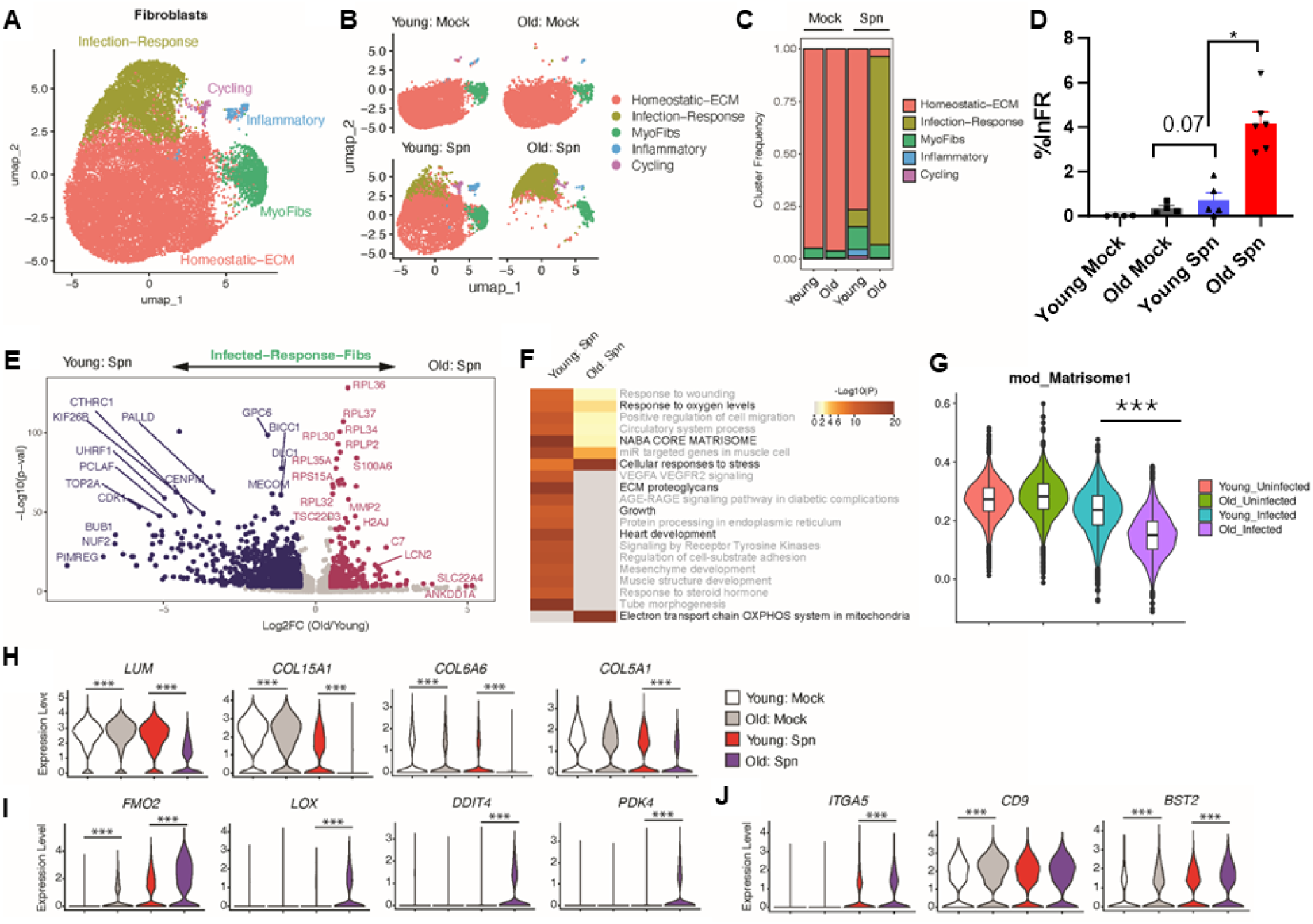
Aging skews cardiac fibroblasts toward inflammatory and maladaptive remodeling phenotypes. **A**) UMAP visualization of cardiac fibroblast subsets at 96-hpi. **B**) UMAP visualization of cardiac fibroblasts in different experimental conditions. **C**) Frequencies of fibroblast subsets across young mock, aged mock, young *Spn*, and aged *Spn* groups. **D**) Identification and quantification of infection responsive fibroblasts (InRF) by flowcytometry. **E**) Differential gene expression analysis of InRFs from young and aged *Spn*-infected hearts shown as a volcano plot. **F**) Pathway enrichment analysis in fibroblasts from young and aged *Spn*-infected mice. G–J) Quantification of fibroblast-associated inflammatory and remodeling programs. **G**) Matrisome module scores across experimental conditions. **H**) Expression of extracellular matrix-synthesizing genes. **I**) Expression of oxidative stress- related genes. **J**) Expression of interferon response-associated genes. P-values compare young and old mice. ***p < 0.001 by Wilcoxon rank-sum test.

Fibroblast composition was strongly shaped by both infection and age. In mock-treated young and aged hearts, fibroblasts predominantly occupied the homeostatic-ECM cluster, consistent with a quiescent, matrix-maintaining phenotype. In contrast, *Spn* infection drove a pronounced expansion of InRFs with broad transcriptional reprogramming, indicating both quantitative and qualitative shifts in fibroblast state (**Fig.5C, D**). Notably, InRFs from aged-infected hearts upregulated ribosomal genes together with inflammatory and stress-responsive factors, including LCN2, S100A6, and MMP2, and TSC22D3 (**Fig.5I**), consistent with a stress-adapted, proinflammatory phenotype.

Pathway analysis revealed a fundamental divergence in fibroblast response programs with age. Fibroblasts from aged mice downregulated reparative pathways, including response to oxygen, NABA core matrisome, ECM proteoglycans, growth, and heart development (**Fig.5F**). while preferentially upregulating pathways linked to cellular stress and mitochondrial metabolism, including oxidative phosphorylation (OXPHOS). Matrisome module scoring demonstrated infection-induced suppression of ECM programs, which was most pronounced in aged fibroblasts (**Fig.5G**). Concordantly, structural ECM genes (LUM, COL15A1, COL6A6, COL5A1) were reduced in aged versus young, infected fibroblasts, indicating impaired matrix remodeling capacity (**Fig.5H**). Instead, aged fibroblasts upregulated oxidative stress and metabolic regulators, such FMO2, LOX, DDIT4, and PDK4 (**Fig.5I**).

Age-dependent rewiring also extended to cell–matrix and inflammatory signaling. ITGA5 was strongly induced in aged fibroblasts upon infection, while CD9 remained constitutively elevated with age, and the interferon-stimulated gene BST2 was induced in both groups but reached higher levels in aged fibroblasts (**Fig.5J**). Flow cytometry confirmed expansion of InRFs defined by a CD9⁺ITGA5⁺BST2⁺ phenotype, validating the transcriptional state shift at the protein level in aged mice (**Fig.5D**). Collectively, these data support a model in which aged fibroblasts fail to mount a coordinated reparative response centered on ECM production and tissue remodeling and instead are redirected towards a stress-dominant, inflammatory, and metabolically reprogrammed state that may compromise effective cardiac repair following infection.

### Aging amplifies LCN2 expression across cardiac cell populations

Given the established association between LCN2 and heart failure^28,29^, we next investigated how aging shapes LCN2 expression in the infected heart. LCN2 was markedly upregulated in neutrophils, macrophages, and fibroblasts from aged *Spn*-infected mice (**Fig.4D, 4I and 5E**). To define its broader cellular sources, we quantified LCN2 expression across all major cardiac cell populations. Interestingly, LCN2 was broadly elevated in aged, infected hearts, with increased expression in endothelial cells, fibroblasts, myeloid cells, mesothelial cells, smooth muscle cells, and adipocytes (**Fig.6A**). Focusing on myeloid and stromal compartments, LCN2 expression was significantly higher in neutrophils, monocyte–macrophage populations, and tissue-resident macrophages from aged versus young infected mice (**Fig. 6B**). Within the fibroblast compartment, InRFs, myofibroblasts, and inflammatory fibroblasts all showed increased LCN2 expression in aged, infected hearts, with InRFs constituting the principal fibroblast source **(Fig. 6C)**. Together, these findings reveal a striking age-associated amplification of the LCN2 axis across immune and stromal compartments, highlighting LCN2 as a potential therapeutic target in pneumonia-associated cardiac dysfunction in aging.

**Fig. 6.**
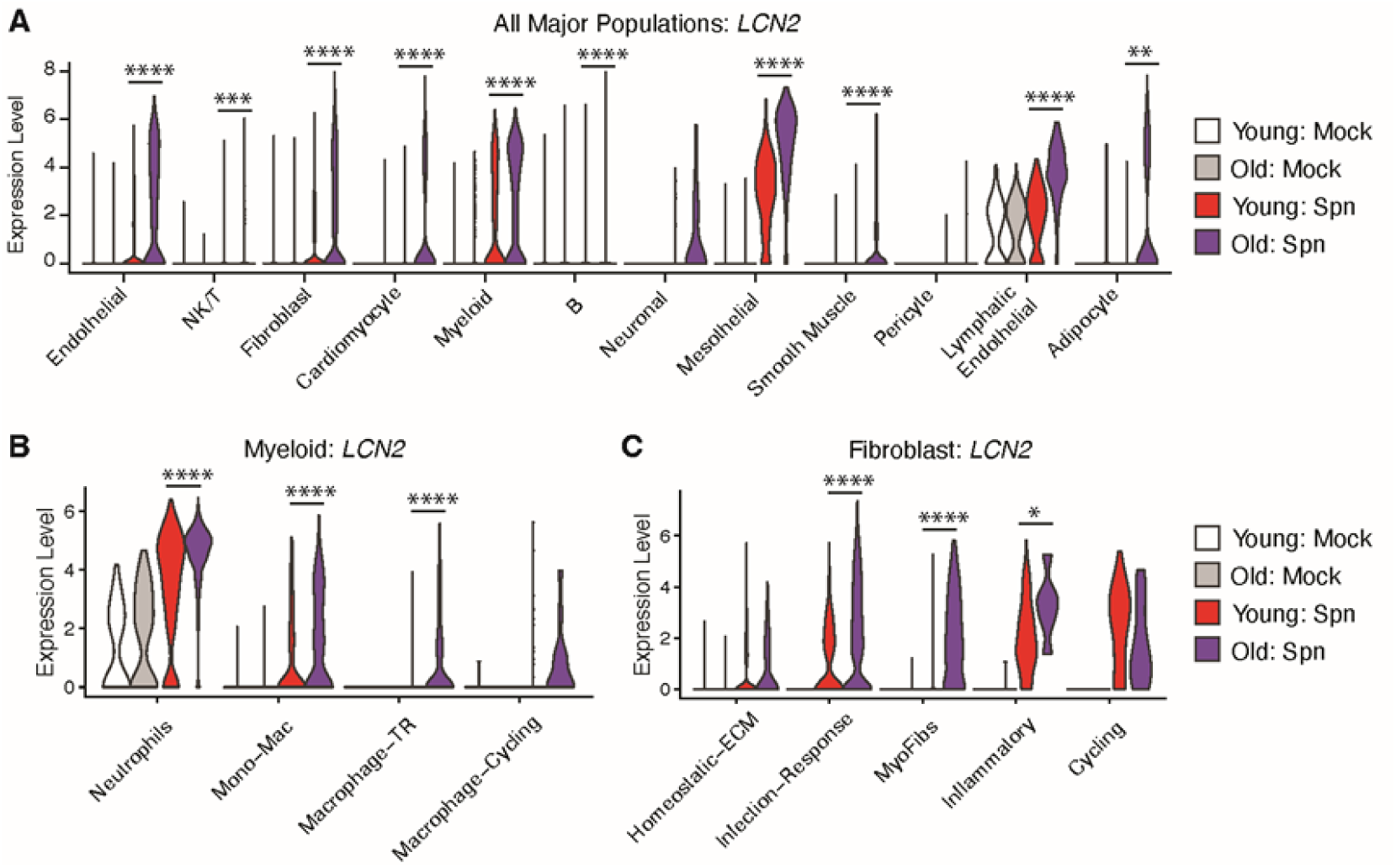
Cell-type specific LCN2 is strongly induced in aged mice. LCN2 expression across different cardiac cell populations at 96-hpi. **A**) LCN2 expression across all cardiac cell types identified by scRNA-seq. **B**) LCN2 expression within myeloid subsets. **C**) LCN2 expressions within fibroblast subsets. P-values compare young and old mice. ***p < 0.001 by Wilcoxon rank-sum test.

### Heightened inflammatory response predisposes aged mice to chronic cardiac dysfunction

Having observed that aging predisposes mice to heightened cardiac inflammation during the convalescent phase of *Spn* infection, we next asked whether early events during the acute phase of infection establish a proinflammatory state in the aged heart. We hypothesized that exaggerated activation of cardiac stromal and immune cells during acute infection drives sustained inflammatory signaling, thereby amplifying responses during recovery. To test this, we quantified chemokines and cytokines in cardiac tissue and serum at 24-hpi. In the heart, TNF-α and IL-1β were modestly elevated in aged *Spn*-infected mice compared with young, infected mice but did not reach statistical significance (**Fig.7A**). Systemically, all measured mediators (TNF-α, IL-1β, CXCL2, IL-6, and S100A8/9) were higher in the serum of aged versus young *Spn*-infected mice, consistent with amplified systemic inflammation (**Fig.7B**).Together, these data indicate that aging predisposes mice to hyperactivation of cardiac and systemic inflammatory responses during *Spn* infection.

**Fig. 7.**
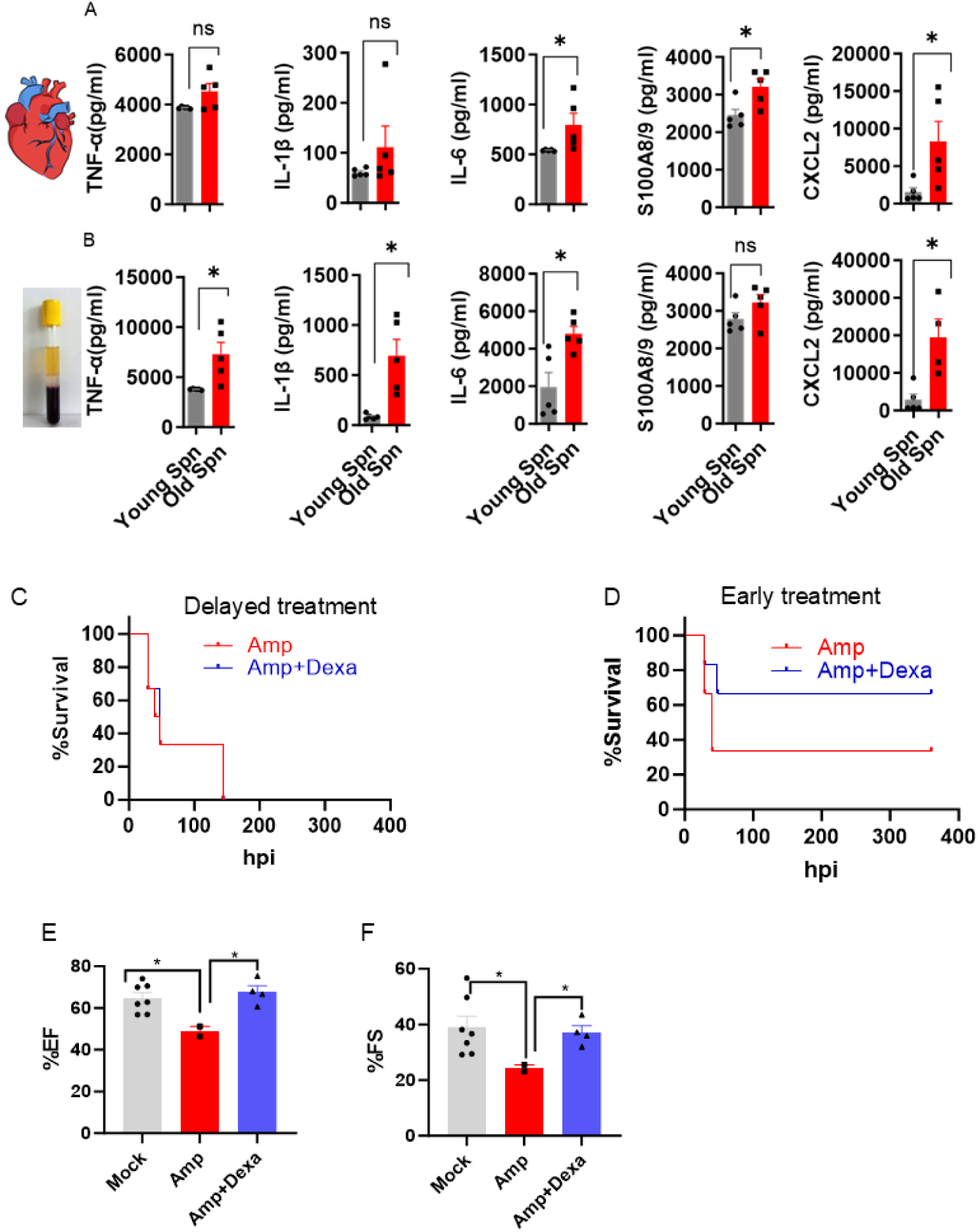
Heightened inflammatory responses predispose aged mice to chronic cardiac dysfunction. Young and aged mice were infected intraperitoneally with *Spn* (0.5 × 10³ CFU), and cardiac and serum cytokines were measured at 24 h post-infection (hpi) (n = 5 per group). Concentrations (pg ml⁻¹) of TNF-α, IL-1β, IL-6, S100A8/9, and CXCL2 in heart homogenates (**A**) and serum (**B**). Effect of early versus delayed antibiotic and dexamethasone treatment in aged mice. **C**) Aged mice were infected and treated with ampicillin (100 mg kg⁻¹) with or without dexamethasone (10 mg kg⁻¹) starting at 21 hpi (delayed treatment; Amp group n = 3, Amp+Dexa group n = 6). **D**) Aged mice treated starting at 18 hpi (early treatment; Amp group n = 6, Amp+Dexa group n = 6). Mice received drugs every 10 h for 3 days, and survival was monitored. Cardiac function at 15 days post-infection, showing ejection fraction (%EF; **E**) and fractional shortening (%FS; **F**). Each dot represents one mouse; bars indicate mean ± SEM. Cytokine/chemokine and treatment experiments were performed once each. Statistical comparisons were performed using Student’s test (*P < 0.05).

Because early inflammation appeared to be a critical determinant of mortality, we conducted a proof-of-concept intervention to test whether early modulation of infection and inflammation could improve survival. Aged mice infected with *Spn* (0.5×10³ CFU) were assigned to early- or delayed-treatment groups and received ampicillin (100 mg/kg body weight) plus dexamethasone (10 mg/kg body weight) beginning at 18-hpi (early group) or 21-hpi (delayed group) followed by additional doses every 10 hours for a total of four treatments. Early intervention resulted in ≥ 60% survival, whereas all mice in the delayed-treatment group succumbed to infection (**Fig.7C**, **D**). These findings indicate that early inflammatory responses are major drivers of mortality in aged hosts and that timely suppression of excessive inflammation can substantially improve outcomes. Consistent with improved recovery, dexamethasone-treated survivors exhibited preserved cardiac function at 15-DPI, with ejection fraction and fractional shortening returning to baseline (**Fig.7E, F**). Collectively, these data suggest that aging creates a temporarily sensitive window of inflammatory hyperactivation during *Spn* infection, in which early containment of inflammation alongside antibiotic therapy is critical to enhance survival and prevent chronic cardiac dysfunction.

### Bone marrow transfer of young immune cells does not rescue aged mice

To test rejuvenation of the hematopoietic compartment can rescue aged hosts during *Spn* infection, we generated bone marrow chimeras by transplanting CD45.1⁺ young donor bone marrow into lethally irradiated aged CD45.2⁺ recipients. Hematopoietic reconstitution was efficient but incomplete, with approximately 80% CD45.1⁺ donor-derived and 20% residual CD45.2⁺ host-derived cells in peripheral blood (**Fig.8A, B**).

**Fig. 8.**
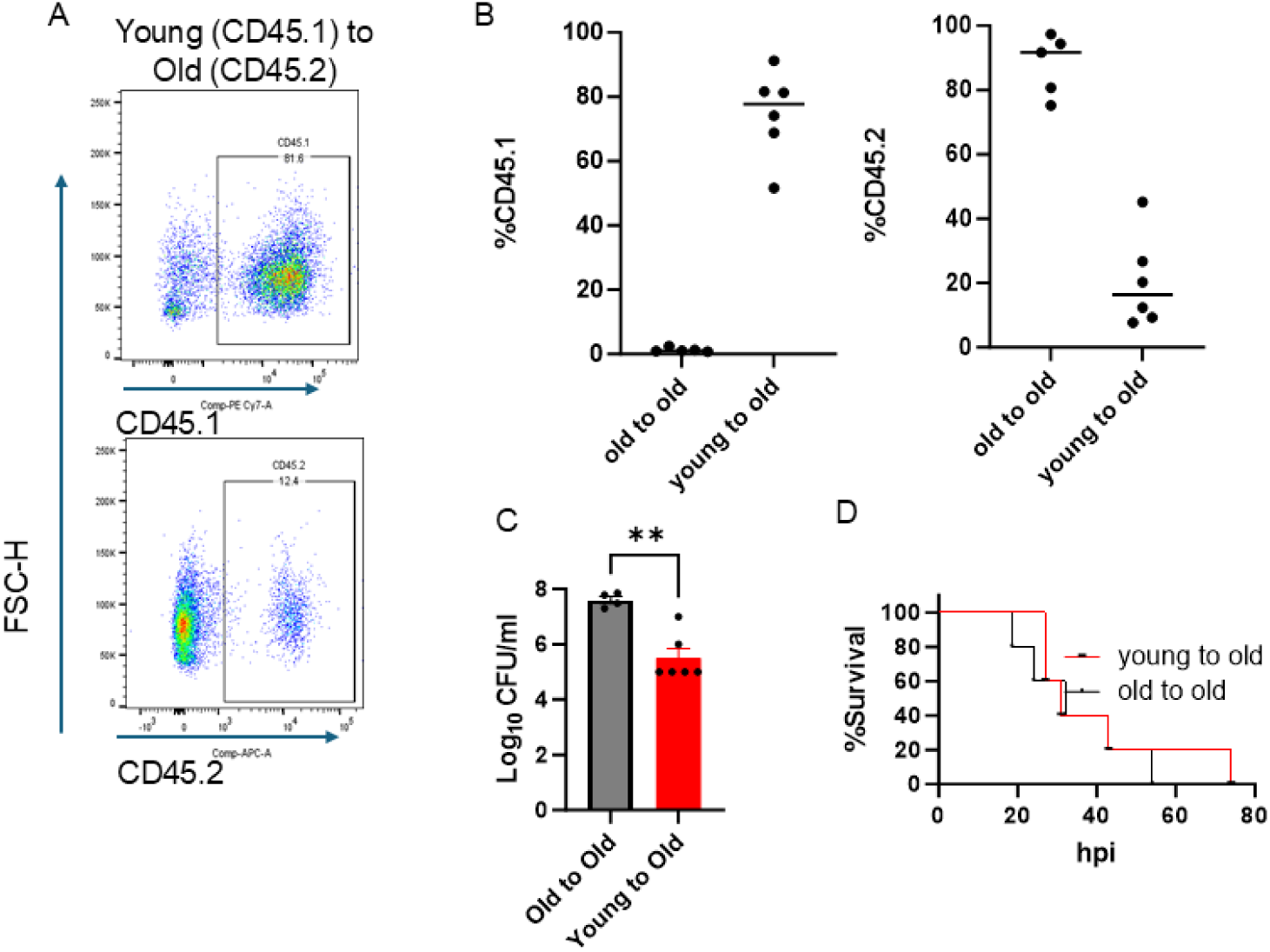
Young bone marrow cells do not provide survival advantage to aged mice. Peripheral blood flow cytometric analysis of aged recipient mice reconstituted with young (CD45.1⁺) or aged (CD45.2⁺) bone marrow, performed 7 weeks after transplantation. **A**) Representative flow cytometry plots showing donor marrow engraftment in recipient mice. **B**) Quantification of CD45.1⁺ and CD45.2⁺ donor-derived cells in peripheral blood. Bacterial burden (**C**) and survival (**D**) of aged recipients reconstituted with young (CD45.1⁺) or aged (CD45.2⁺) bone marrow following *Spn* infection at 8 weeks post-transplant. Each dot represents one mouse; bars indicate mean ± SEM. The experiment was performed once. Statistical comparisons were performed using unpaired two-tailed Student’s t-test (**P<0.01).

Chimeric mice were then infected with *Spn* and treated with antibiotics starting at 18 hours post-infection (hpi) for a total of four doses. Although young to old chimeras displayed reduced bacteremia compared with aged to aged (**Fig. 8C**), this improvement in bacterial control did not confer a survival advantage, as all mice in both groups succumbed before 80 hpi (**Fig.8D**). These findings indicate that hematopoietic rejuvenation alone is insufficient to rescue mortality in aged hosts. Instead, they suggest that non-hematopoietic, tissue-resident, or systemic aging-associated factors, potentially within the cardiac or stromal compartments, play a critical role in driving poor outcomes during pneumococcal infection in aged mice.

## Discussion

Aging is a major risk factor for severe pneumococcal pneumonia and PA-MACE^8,30^. Here, we show that aged mice exhibit markedly increased susceptibility to pneumococcal infection, characterized by higher bacteremia, reduced survival, exacerbated cardiac dysfunction, and dysregulated immune and stromal responses. Our data reveal distinct age-dependent defects in early antimicrobial defense coupled to persistent inflammatory and tissue-remodeling programs that likely sustain inflammation and cardiac dysfunction during acute and convalescent phases of pneumococcal infections.

Aged mice remained highly susceptible to *Spn* infection even at substantially reduced inocula, and echocardiography demonstrated pronounced systolic dysfunction with reduced heart rate, stroke volume, ejection fraction, and cardiac output, consistent with decompensated heart failure **(Fig.1**, **Fig.2)**. These observations align with clinical data showing that CAP is a major cause of morbidity and mortality in older adults and confers a long-term risk of cardiac dysfunction^31–33^. Although pneumolysin is known to directly injure cardiomyocytes³, our findings indicate that an aged cardiac environment, shaped by both inflammaging and infection-induced responses, plays a critical role in amplifying cardiac injury.

Consistent with heightened cytokine and chemokine production (TNF-α, IL-1β, IL-6, S100A8/9, and CXCL2) (**Fig.7A, B**), aged *Spn*-infected mice displayed increased neutrophil accumulation in the heart and elevated Ly6G expression on circulating neutrophils, indicative of enhanced recruitment and activation^34^ (**Fig.4G, H**). This is in line with reports of increased neutrophil frequencies in elderly patients with pneumococcal pneumonia and supports the concept that aging remodels neutrophil biology and is associated with multiple functional defects^34–38^. ScRNA-seq analysis revealed a hyperinflammatory neutrophil state in age infected hearts, with upregulation of S100A8/9 and LCN2 but downregulation of pathways involved in endocytosis and degranulation. Strikingly, elevated CXCL2 in aged hearts, together with reduced CSF3R expression in aged neutrophils **(Fig.7A**, **Fig.4D)**, suggests an altered signaling milieu that may impair that balance between activation, survival and function^39^. Furthermore, reduced expression of lysosome-associated genes such as CTSB, LAMP2, and CTSD **(Fig.4F)** indicates defective lysosome-dependent antimicrobial activity^40–42^. Together, these data support a model in which aging enhances neutrophil recruitment yet rewires effector programs towards a destructive inflammatory phenotype with diminished antimicrobial capacity, thereby increasing cardiac vulnerability and identifying neutrophil inflammatory pathways as potential therapeutic targets.

Macrophages, which are essential for bacterial clearance, efferocytosis, and maintenance of electrical and inflammatory homeostasis^5,24,43^, were reduced in number and showed profound transcriptional reprogramming in aged, infected hearts **(Fig.4C)**. Aged macrophages upregulated inflammatory and stress-related genes, including TREM1, CCL17, CXCL13, IL1R1, GSR, and ITGAM **(Fig.4I)**, consistent with heightened inflammatory polarization and leukocyte recruitment. While macrophages play critical role in engulfing infected, aged and apoptotic neutrophils^44,45^. At the same time, genes central to efferocytosis and tissue repair—GAS6, AXL, DOCK1, and related pathways, were downregulated **(Fig.4I-K)**, indicating impaired clearance of apoptotic cells and defective resolution. Given the central role of GAS6–AXL signaling in recognizing of apoptotic neutrophils and DOCK1 in cytoskeletal rearrangements required for engulfment, this signature points to a severe efferocytosis defects in aged cardiac macrophages^46–48^. Thus, aging appears to disrupt macrophage resolution programs via impairment of the GAS6–AXL axis, leading to persistence of activated neutrophils, ongoing inflammation, and inadequate repair.

Beyond immune cells, our single-cell analysis revealed that aging profoundly reshapes fibroblast behavior in the infected hearts. *Spn* infection induced an activated fibroblast program that was selectively amplified in aged mice, driven by expansion of InRFs with inflammatory and interferon-responsive signatures, including LCN2, S100A6, and BST2, suggest that the aged cardiac stroma responds to bacterial challenge with a heightened and maladaptive transcriptional program. These InRFs, defined by CD9⁺ITGA5^high^BST2^high^ expression and elevated ITGA5, LOX, and BST2 transcripts **(Fig.5D, E, J)**, share features with inflammatory fibroblasts states described in lung injury and fibrosis models^49^. Matrisome module scoring and gene expression analyses revealed infection-induced suppression of ECM programs, particularly in aged fibroblasts, with downregulation of structural ECM genes (LUM, COL15A1, COL6A6, COL5A1) and concomitant upregulation of oxidative stress and metabolic regulators such as FMO2, LOX, DDIT4, and PDK **(Fig.5G-I)**. These changes point to a shift away from coordinated reparative ECM remodeling toward a stress-dominant, metabolically reprogrammed phenotype. LCN2 was also robustly expressed by aged fibroblasts, further implicating fibroblast-derived LCN2 in cardiac dysfunction^28,29^. Collectively, these findings suggest that fibroblasts are not passive bystanders but central drivers of age-dependent cardiac vulnerability, and that stromal–immune crosstalk forms a key mechanistic link between aging, inflammation, and infection-induced cardiac dysfunction.

In parallel with the exaggerated inflammatory response during convalescence, aged, infected mice mounted a markedly heightened inflammatory response during the acute phase of infection, both in the heart and systemically. TNF-α, IL-1β, IL-6, and S100A8/9, CXCL2 were substantially elevated in aged hearts and serum **(Fig.7A, B)**, mediators known to promote neutrophil recruitment, amplification of cytokine production, and cardiac stress^36,37^. This heightened early inflammatory milieu likely contributes to the severe impairment of cardiac performance observed in aged *Spn*- infected mice. Our findings further demonstrate that early inflammatory containment can restore cardiac function and improve survival **(Fig.7C-F)**, highlighting immune–stromal inflammatory circuits as therapeutically actionable targets in elderly individuals with severe pneumococcal disease. Although young hematopoietic cells improved bacterial clearance in aged recipients, they failed to confer any survival benefit **(Fig.8A-D)**. These bone marrow chimera results indicate that cardiac dysfunction and death in aged mice are not primarily driven by bacterial burden or hematopoietic insufficiency alone. Rather, they suggest that non-hematopoietic, tissue-resident determinants may be disease drivers aged hearts. One additional possibility is that cardiac-resident immune cells are incompletely replaced by young cells or remain functionally imprinted by the aged microenvironment, or alternatively that the aged cardiac niche actively reprograms infiltrating young immune cells toward an aged-like inflammatory state. These possibilities cannot be fully resolved in the present study and warrant further investigation in future work.

Together, our findings support a model in which aging exacerbates pneumococcal-induced cardiac dysfunction and inflammation through convergent mechanisms: (1) amplified neutrophil, and macrophage-driven inflammation with impaired macrophage efferocytosis, and (2) age, and infection-induced fibroblast activation with defective ECM remodeling and stress-dominant reprogramming. These processes likely act synergistically to sustain cardiac inflammation during convalescence and impede resolution of tissue injury. By redefining how age shapes host, pathogen interactions at the level of immune and stromal circuits, this work reveals cellular pathways that may underlie the heightened vulnerability of older individuals to PA-MACE and other infection-associated cardiac complications. Future studies should focus on strategies to enhance macrophage-mediated clearance and resolution in aged hosts and to attenuate both immune and stromal cells, driven cardiac inflammation in older adults.

## Materials and methods

### Ethics statement

All animal procedures in this study were approved by The Ohio State University Institutional Animal Care and Use Committee (IACUC) under protocol 2020A00000004-R1.

### Mice

C57BL/6 mice 10-12 weeks (young) or 18-20 months (aged) were purchased from Charles River Laboratories (Wilmington, MA) via a contract with National Institute on Aging. Mice were housed 3–5 per microisolator cage, acclimated for 1 week, and maintained on a 12-h light/dark cycle with ad libitum access to food and water.

### Bacterial growth, murine infection and antibiotic treatment

*Streptococcus pneumoniae* strain TIGR was cultured to the exponential phase in Todd-Hewitt Broth at 37 °C under 5% CO₂ until an optical density at 621 nm of 0.5 was reached^50^. For mutants^51^, spectinomycin (100 µg/mL) or erythromycin (1 µg/mL) was added to the growth medium to maintain selection of the Δply and ΔspxB mutant strains, respectively. All *Spn* strains were generous gifts from Dr. Carlos J Orihuela (Department of Microbiology, The University of Alabama at Birmingham, Birmingham, AL, USA). Young (10-week-old) and aged (18–20-month-old) female C57BL/6 mice were infected intraperitoneally with 0.5×10³ CFU or 1×10^6^ CFU via intranasal route or otherwise stated. Exponentially grown *Spn* was suspended in 200 µL of sterile saline. Mock-infected control mice received 200 µL of saline. For the low-inoculum experiment, aged mice were infected with a 20-fold lower dose (0.025×10³ CFU), while young mice received the standard 0.5×10³ CFU/mouse. Starting at 18- or 21- or 28-hpi, infected mice were administered filter-sterilized ampicillin (100 mg/kg) dissolved in saline, delivered intraperitoneally in a volume of 200 µL, and this treatment was repeated every 10 hours for 3 days.

### Blood collection and Bacterial Burden assay

Blood was collected from the lateral facial vein of adult mice as previously described^52^. Mice were manually restrained and blood was drawn using a sterile 25–27 G lancet into microcentrifuge tubes. Gentle pressure was applied to achieve hemostasis before returning mice to their home cages. 10 µl blood was immediately serially diluted in PBS and plated on sheep blood agar plate to determine bacterial load.

### Echocardiography

Cardiac function was assessed in vivo using 2D echocardiography (Vevo 2100, VisualSonics) in *Spn*-infected or mock mice at 24 hours post-infection. Mice were anesthetized in an induction chamber with 2% isoflurane in oxygen (1.0 L/min) and placed supine on a heated stage. Chest hair was removed using a depilatory lotion, and anesthesia was maintained at 1.5% isoflurane throughout the procedure. Heart rate was continuously monitored to ensure appropriate anesthetic depth. An MS-400 transducer was used to obtain images along the parasternal long axis (PSLX). M-mode images were recorded at the level of the papillary muscles. Cardiac parameters, including beats per minute, ejection fraction, fractional shortening, and cardiac output were analyzed to assess cardiac function.

### Alveolar macrophage isolation and infection

Young and aged mice were euthanized using CO₂ in accordance with an OSU IACUC-approved protocol. Alveolar macrophages (AMs) were collected by bronchoalveolar lavage (BAL). Lungs were lavaged ten times with 0.5 mL sterile, and endotoxin-free saline (0.9% NaCl). AMs were isolated by centrifugation of the BAL fluid at 600 × g for 5 min. For each experiment, AMs from 3-5 mice were pooled and plated on a 48-well plate. Macrophages were allowed to attach to the plate for 3 hours and were washed with RPMI to remove unbound cells. Cells were resuspended in RPMI containing 5% FBS and allowed to rest overnight. Next day macrophages were infected with pre-opsonized *Spn* (mid-log phase *Spn* incubated with 3% normal mouse serum for 30 min at 37°C with gentle shaking) for 1 h at 10 MOI in RPMI. Plates were centrifuged at 600 rpm for 3 min at room temperature for synchronized infection and further incubated at 37°C for 30 minutes in a CO2 incubator. Cells were washed 3 times with RPMI to remove unbound bacteria. To kill remaining extracellular bacteria, cultures were treated with Geneticin (200 μg/mL) for 30 min at 37°C, followed by three additional RPMI washes. Cells were then incubated for 3 h at 37°C in a CO₂ incubator. At the experimental endpoint, supernatants were removed and intracellular bacteria were released by treating cells with 0.1% Triton X-100 for 5 min on a shaker at room temperature. Colony-forming units (CFU) were quantified by plating serial dilutions of lysates on sheep blood agar plates.

### ELISA

TNF-α (DY410), CXCL2 (DY452), IL-1β (DY401), IL-6 (DY406), and S100A8/A9 (DY8596) levels in serum and cardiac tissue homogenates were quantified using ELISA kits (R&D Systems) according to the manufacturer’s instructions.

### Multielectrode array (MEA) assay

Human induced pluripotent stem cell-derived cardiomyocytes (hiPSC-CMs) were plated as previously described^53^. Briefly, cells were cultured on 0.1% gelatin–coated tissue culture plates using plating medium (FUJIFILM Cellular Dynamics, Inc., WI, USA) and incubated for 48 h in a humidified chamber at 37 °C with 5% CO₂. HiPSC-CMs were subsequently harvested and re-plated (3.0 × 10⁴ cells per well) into fibronectin-coated (50 µg/mL) 24-well CytoView MEA plates (Axion BioSystems, GA, USA). HiPSC-CMs were infected with *Spn* (MOI = 1). Recordings were collected for 5 min at 15 min intervals for up to 48 hours. Data were acquired using AxIS Navigator™ (v2.0.4.21) and analyzed with the Cardiac Analysis Tool™ (v3.1.8) and AxIS Metric Plotting Tool (v2.3.1).

### Cardiac cell isolation

Shortly after euthanasia, mice were perfused with ice-cold PBS until the liver appeared pale yellow. Hearts were then harvested and enzymatically digested using the Multi Tissue Dissociation Kit and gentleMACS Dissociator (Miltenyi Biotec, Auburn, CA, USA) according to the manufacturer’s instructions. The resulting homogenized tissue was suspended in 7.5 mL of RPMI containing 20% FBS and passed through a 100 µm cell strainer. Cells were pelleted by centrifugation at 600 × g and further processed with Debris Removal Solution (Miltenyi Biotec) to remove dead cells and debris. The final cell suspension was resuspended in PBS and used for downstream applications, including single-cell RNA sequencing (scRNA-seq) or flow cytometry.

### scRNA-seq Analysis

Cells were processed according to the Chromium Next GEM Single Cell 3’ Kit v3.1 from 10x genomics. A total of 10,000 cells were processed per sample via standard 10X protocol. Sequencing was performed on the Illumina NovaSeq PE150 targeting 20,000 reads per cell. Using the 10x cloud system, data were aligned to the GRCm38/mm10 transcriptome using the Cell Ranger Count v7.1.0 pipeline.

For downstream analysis, we used Seurat v5.2.0 package to create a Seurat object and used Azimuth-heart for reference annotation. For quality control (QC) and filtering, we visualized QC metrics generated by Seurat analysis, keeping cells between 2,500 and 200 feature counts, as well as cells that have more than 5% mitochondrial counts. Principal component analysis (PCA) was performed, and the default top 2000 differential genes were used for analysis. Uniform Manifold Approximation and Projection (UMAP) and t-Distributed Stochastic Neighbor Embedding (t-SNE) dimensional reductions were used for visualization. Figures were generated using ShinyCell V2.1^54^ and Seurat V5.2.0.

### Flow cytometry analysis

Single-cell suspensions were first stained with Live/Dead Zombie Aqua dye for 10 minutes at room temperature. Cells were then washed twice with FACS buffer (2% FBS, 2mM EDTA, 0.1% sodium azide in PBS) and incubated with mouse Fc Block (Bio-Rad) for 15 minutes to prevent nonspecific antibody binding. Following Fc blocking, cells were stained with a panel of fluorochrome-conjugated (Supplementary Table 1) antibodies targeting relevant cell surface markers. Stained cells were analyzed and quantified using a BD Symphony flow cytometer. Data were processed and analyzed using FlowJo software (FlowJo LLC, OR, USA). Flow cytometric analysis of 30 µl of heparinized whole blood from facial was performed as previously described^55^. Briefly, whole blood was stained for CD45, CD11b, and Ly6G (Supplementary Table 1) antibodies in the dark on ice for 30 minutes. Cells were then washed twice with cold FACS buffer. Red blood cells were lysed, and the samples were washed twice more with ice-cold FACS buffer. Stained cells were analyzed as described above. Similarly, whole blood Flow cytometry for bone marrow chimera was performed with CD45.2 or CD45.1 antibodies. Flow cytometry gating strategies for neutrophils, macrophages, and fibroblasts **(Supplementary Fig. 7)**.

### Bone marrow chimera

Fresh bone marrow cells CD45.1 (young) or CD45.2 (old) were transplanted via single intravenous injection into irradiated (4.5 Gy) female C57BL/6 old CD45.2 or old CD45.2 recipient mice with 5 × 10^6^ whole BM competitor cells. Peripheral blood analysis was performed at 7 weeks and *Spn* infection was given at 8 weeks post bone marrow transplant.

### Statistical analysis

Statistical comparisons were performed using log-rank tests and Student’s T-test. The results are shown as the Mean ± SEM. The threshold for significance was set at p< 0.05.

---

**Supplementary Fig. 1.**
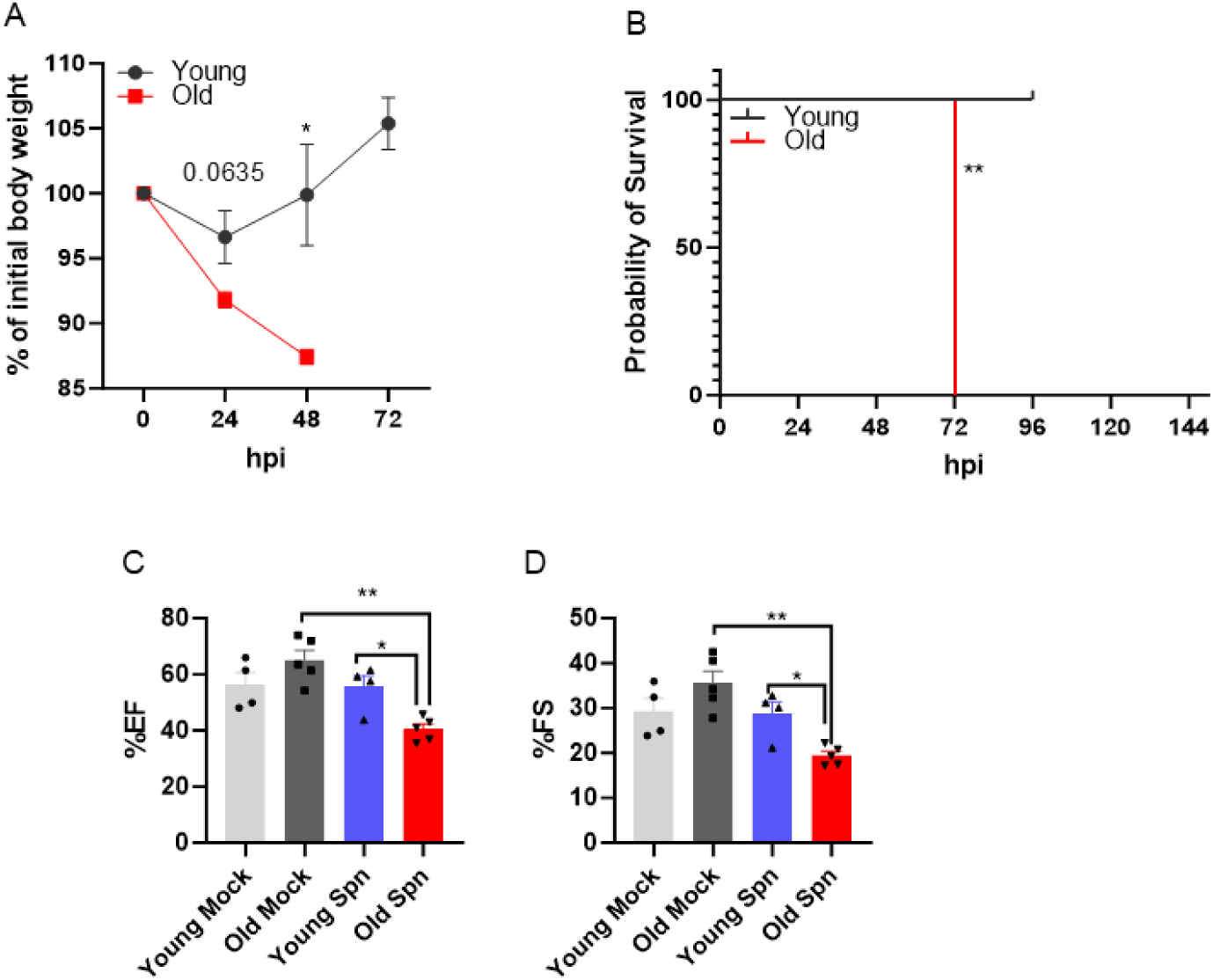
**Intranasal route of infection causes similar cardiac dysfunction in aged mice**. Young and aged mice were infected intranasally with Spn (10^6^ CFU/mice) and assessed for weight changes, survival and cardiac outcomes. A–B) Weight loss and survival curve. C) %EF at 48-hpi. D) %FS at 48-hpi. Experiment was performed once. Statistical significance was determined by using two-tailed Student’s t-test (*P < 0.05, **P<0.01).

**Supplementary Fig. 2.**
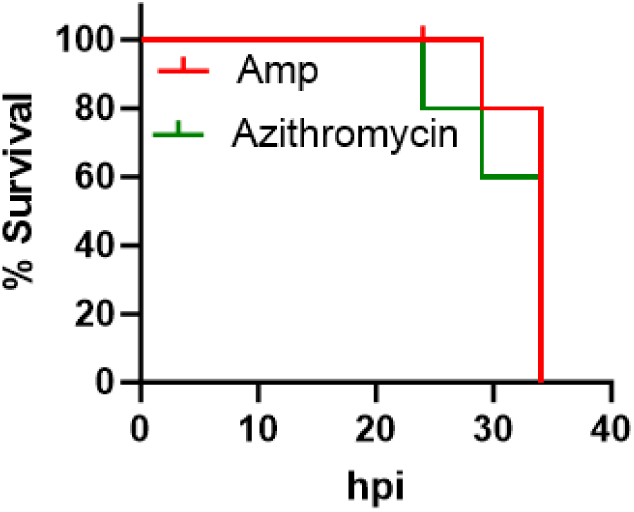
Bacteriostatic antibodies failed to improve survival in aged mice compared with lytic antibodies. Aged mice were given ampicillin or azithromycin 100 mg/kg body weight, dosage as stated in Figure 1 (Ampicillin group n=10, Azithromycin group n=9). Experiment was performed once.

**Supplementary Fig. 3.**
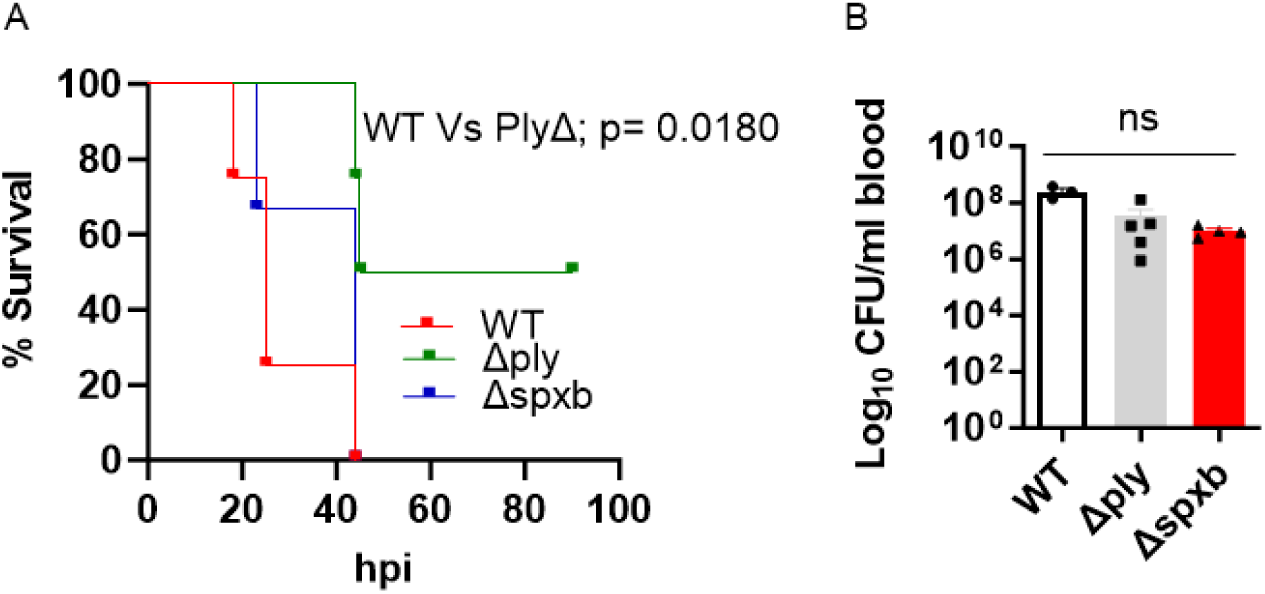
Pneumolysin is required for lethality in aged mice. A-B) Mice were infected with wild-type *Spn* or Δply or ΔspxB with 0.5×10^3^ CFU, intraperitoneal; given antibiotics as stated in Figure 1 (wild type *Spn* n= 4, Δply n=4 ΔspxB n= 5). A) Survival curves. B) Bacterial burden in blood at 24-hpi. Experiment was performed once. Statistical significance was determined by using students t-test; ns = non-significant P value.

**Supplementary Fig. 4.**
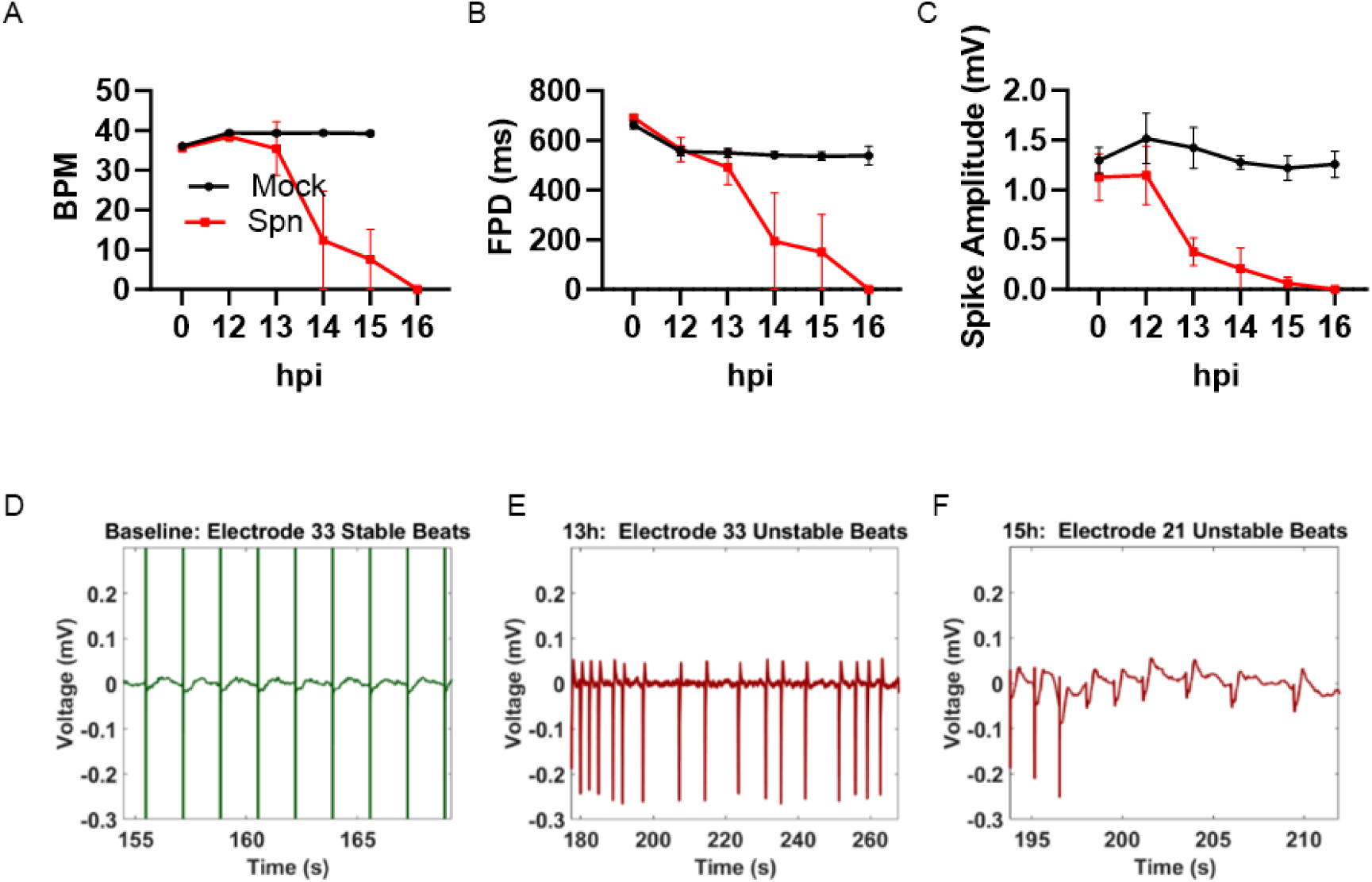
Streptococcus pneumoniae infection disrupts cardiomyocyte electrical function. A-E) hiPSC-CMs were mock or infected with *Spn* (M.O.I = 5, mock n=4, *Spn* n=3). A–C) Multi-electrode array (MEA) measurements showing beating rate (BPM), field potential duration (FPD), and spike amplitude of cardiomyocytes over time following mock treatment or infection with *Spn* D–F) Representative MEA voltage traces (D) Baseline cardiomyocyte (E) at 13-hpi (F) by 15-hpi. Experiment was performed once.

**Supplementary Fig. 5.**
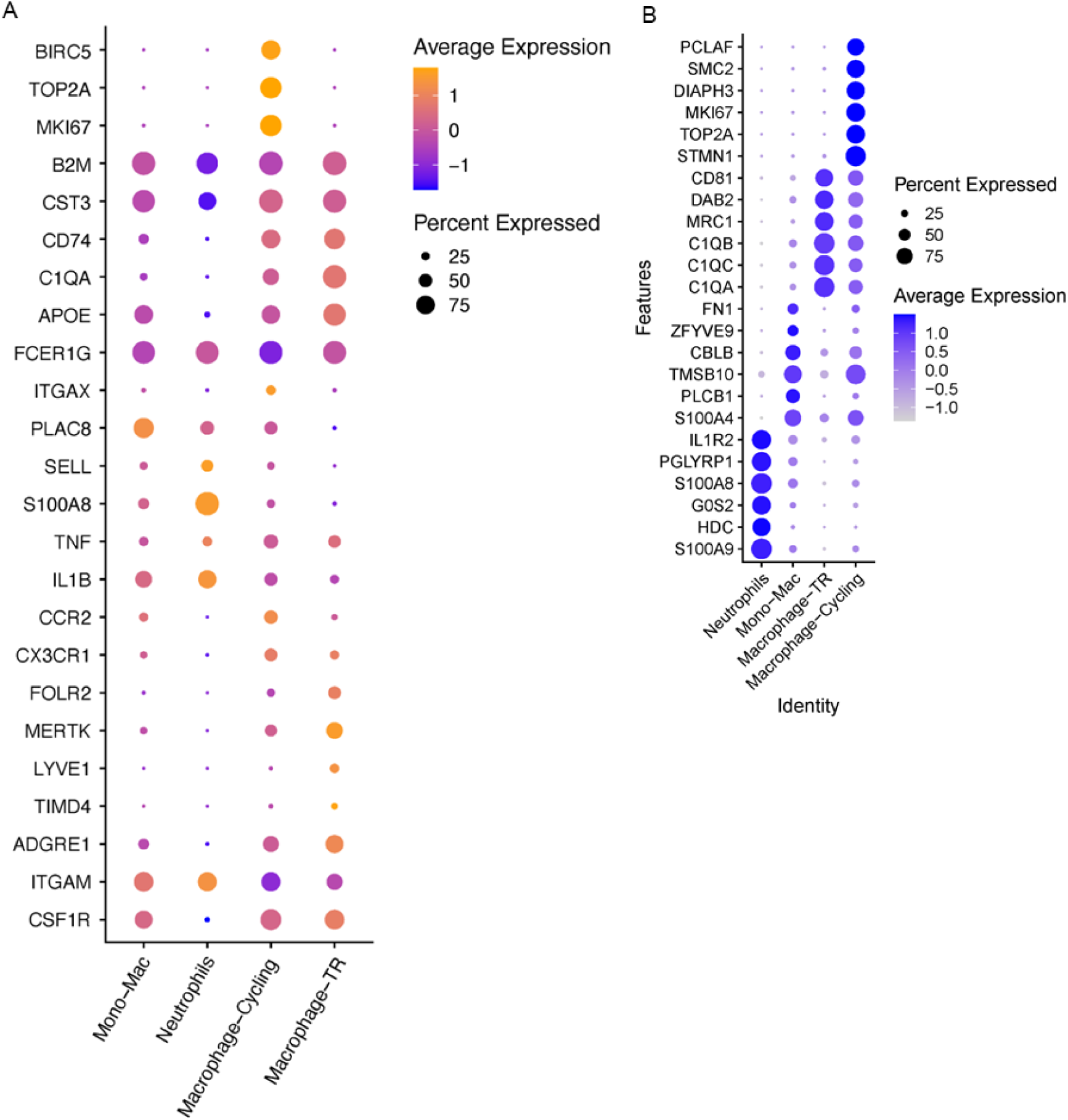
**Identification of myeloid cell subsets**

**Supplementary Fig. 6.**
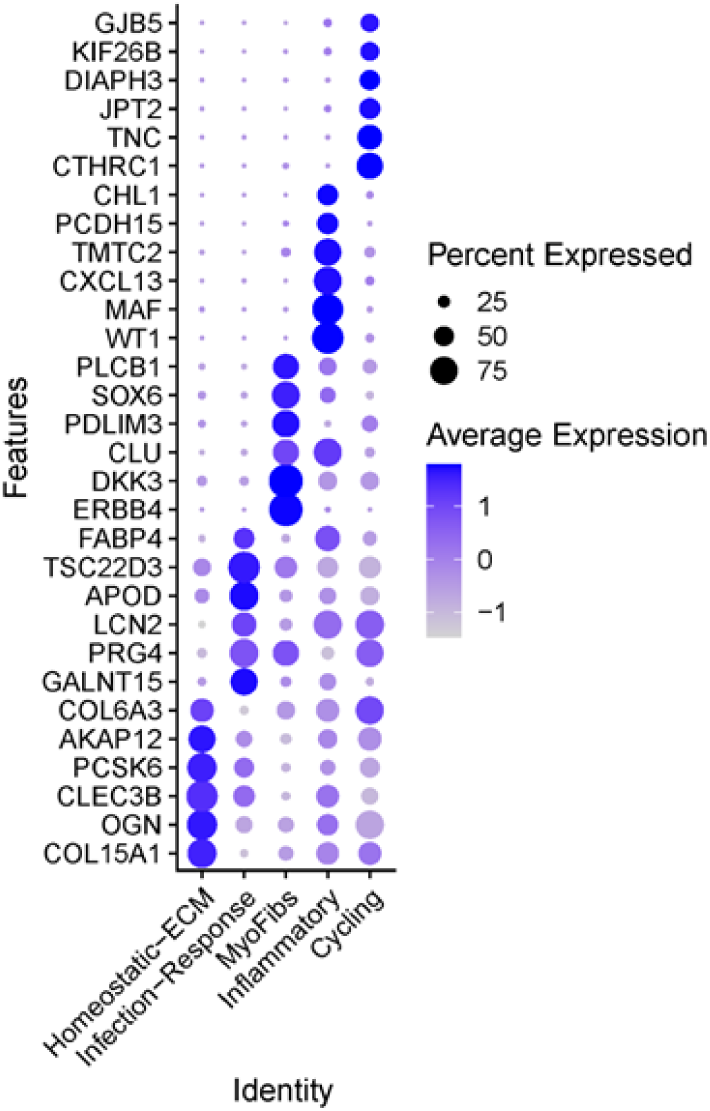
**Identification of fibroblast subsets**

**Supplementary Fig. 7.**
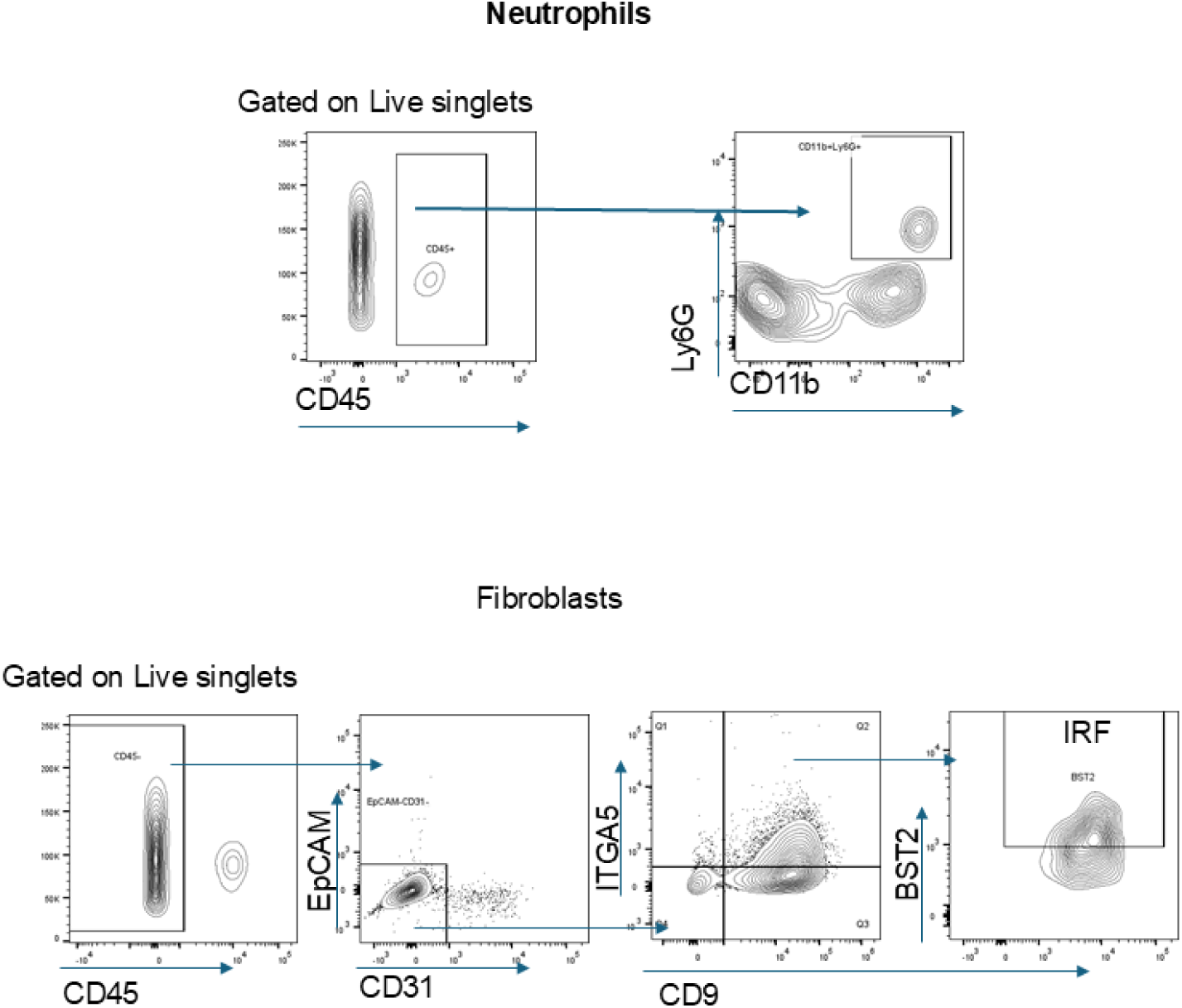
**Flow cytometry gating strategies for neutrophils, macrophages, and fibroblasts**

